# Diverse Epithelial Cell Populations Contribute to Regeneration of Secretory Units in Injured Salivary Glands

**DOI:** 10.1101/2020.06.29.177733

**Authors:** Ninche Ninche, Mingyu Kwak, Soosan Ghazizadeh

## Abstract

Salivary glands exert exocrine secretory function to provide saliva for lubrication and protection of the oral cavity. Its epithelium consists of several differentiated cell types including acinar, ductal and myoepithelial cells that are maintained in a lineage-restricted manner during homeostasis or after mild injuries. Glandular regeneration following a near complete loss of secretory cells, however, may involve cellular plasticity, although the mechanism and extent of such plasticity remain unclear. Here, by combining lineage-tracing experiments with a model of severe glandular injury in the mouse submandibular gland, we show that *de novo* formation of secretory units involves induction of cellular plasticity in multiple non-acinar cell populations. Fate-mapping analysis revealed that although ductal stem cells marked by cytokeratin K14 and Axin2 undergo a multipotency switch, they do not make a significant contribution to acinar regeneration. Intriguingly, more than 80% of regenerated acini derive from differentiated cells including myoepithelial and ductal cells that dedifferentiate to a progenitor-like state before redifferentiation to acinar cells. The potential of diverse cell populations serving as a reserve source for acini widens the therapeutic options for hyposalivation.

**Summary:** Salivary glands rely in recruitment of committed and fully differentiated cell populations as well as stem cells to ensure rapid regeneration and recovery of secretory cells.

## INTRODUCTION

The epithelial tissue in salivary glands is a slowly renewing tissue, organized into a highly branched tubular structure composed of saliva secreting terminal units or acini, a ductal network of intercalated, granular (in rodents), striated and excretory ducts for modifying and transporting saliva, and a contractile myoepithelium to aid in expulsion of saliva from acini through ducts (Redman, 1994, Amano et al., 2012). In the mouse submandibular gland (SMG), which is the most complex among salivary glands, acinar, myoepithelial and granular duct (GD) cells are considered as highly-specialized and well-differentiated cell populations that could be easily distinguished by their unique morphology and expression of cell-specific biomarkers (Fig. 1A). The intercalated duct (ID) that connects several acini to a GD, is the smallest epithelial compartment composed of two distinct subsets of cuboidal cells with a relatively undifferentiated morphology (Denny et al., 1997). These cells are distinguished based on the expression of cKit and cytokeratin K14 which are known to mark multipotent embryonic progenitors during salivary gland development (Kwak et al., 2016, Lombaert et al., 2013). However, cell proliferation in the adult SMG is not limited to ID cells as well-differentiated cells in various epithelial compartments undergo mitosis, thus contributing to the slow cellular turnover under steady-state conditions (Denny et al., 1997, Kwak and Ghazizadeh, 2015). Furthermore, recent genetic lineage tracing studies have shown that various cell types in the mature SMG are maintained in a lineage-restricted manner, either by contribution from unipotent stem/progenitor cells or by self-duplication of differentiated cells (Kwak et al., 2016, Kwak et al., 2018, Aure et al., 2015, Emmerson et al., 2017, May et al., 2018). To date, only a single cell population in the SMG has been discovered that displays characteristics assigned to tissue stem cells, including the capacity for long-term clonal self-renewal and for generation of at least one differentiated cell progeny during homeostasis (Kwak et al., 2016, Clevers and Watt, 2018). This cell population was initially identified as a small subset of ductal cells residing at the ID/GD junction and expressing the basal cell markers K14 and K5 but not α-smooth muscle actin (SMA) which is specific to myoepithelial cells (MECs) or cKit which marks ID cells (Fig. 1A)(Kwak et al., 2016). Genetic labeling and long-term lineage tracing studies showed that K14-expressing ductal cells function as an actively cycling, long-term self-renewing cell population that generates a progeny destined to differentiate into K19^+^ ductal cells in the GD (Kwak et al., 2016). Subsequence studies using other basal cell-specific markers including cytokeratin K5 and transcription factor TP63, or Axin2, a target of WNT signaling, have confirmed these initial findings (Weng et al., 2018, May et al., 2018, Song et al., 2018). Moreover, *ex vivo* studies have verified the regenerative capacity of K14-expressing cells in organoid cultures and upon intra-glandular transplantation which are often used to assess stem cell function (Kwak et al., 2018, Xiao et al., 2014). On the other hand, although cKit has long been considered as a marker for the adult salivary gland tissue stem cells (Hisatomi et al., 2004, Lombaert et al., 2008), recent *in vivo* and *ex vivo* genetic labeling and lineage tracing studies have not been supportive of this hypothesis (Kwak et al., 2018).

**Figure 1.**
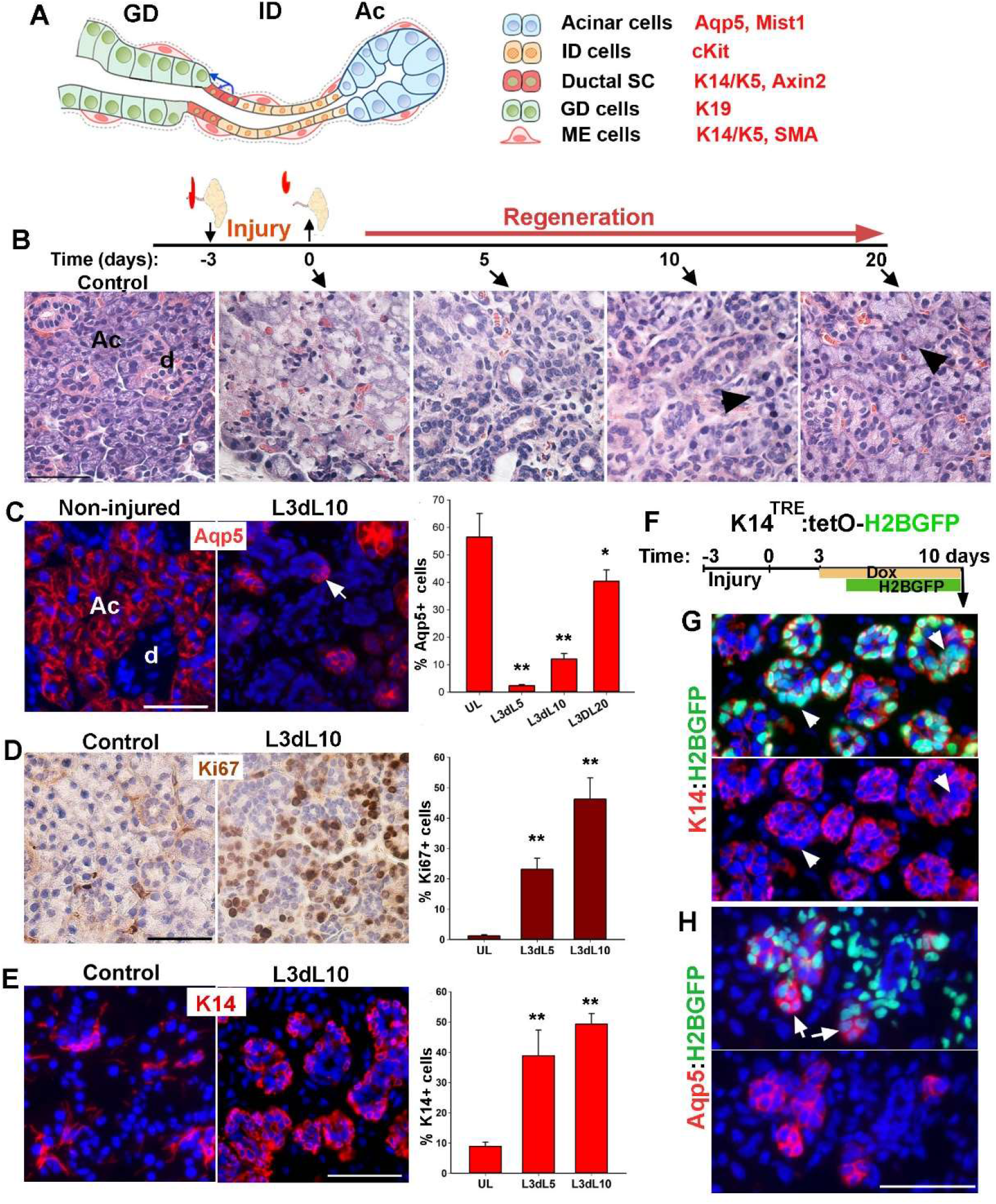
Characterization of salivary gland regeneration in a model of severe injury. (A) Schematic representation of secretory complex in mouse SMG and specific markers used to identify each cell type. Ac is acini, ID is intercalated duct and GD is granular duct. (B) Time-line of duct ligation/deligation injury and images of H&E stained sections of SMG at the indicated time points. Arrowheads point to acini. (C) Distribution and quantification of Aqp5^+^ cells in control (UL) and injured gland collected at 5-, 10- and 20-days post-deligation (L3dL). Arrow points to proacinar cells budding from tubular structures. (D-E) Distribution and quantification of Ki67 proliferation marker and K14 at days 5 and 10 post ligation when tubular structures are present. Image shown is from day 10 post-deligation (L3dL10). Data are mean+s.e.m (n=3 females), one way ANOVA test, *P<0.05 and ** P<0.001. (F) Timeline of nuclear tracking of K14-H2BGFP, (G-H) Images of regenerative gland stained for K14 or Aqp5. Arrowhead in G points to nuclear GFP in K14^neg^ cells. Arrows point to acini-like structures with dim green fluorescence in their nuclei. Images are representative of n=3 female mice. Scale bar=50 μm.

Cell behavior and mechanisms maintaining cell lineages during homeostasis often change dramatically in response to injury (Merrell and Stanger, 2016). Like many other tissues, salivary glands have a capacity to regenerate even after substantial damage, which may involve de-differentiation of various cell types (reviewed in (Denny et al., 1997). Two classic injury models in rodents have been used extensively to study functional recovery and regeneration in salivary glands. In these models, injury is induced either by exposure of the neck to ionizing radiation or by unilateral ligation of the main excretory duct of salivary glands. Irradiation of the gland with a moderate dose (15 Gy) of radiation induces a bi-phasic injury with the initial cell loss being compensated by rapid cell proliferation within 10 days, followed by extensive infiltration of inflammatory cells and degeneration of acini during the late phase of injury (>90 days) (Urek et al., 2005, Bralic et al., 2005). Lineage tracing of ductal and acinar cells in this model of injury have shown that recovery after the early phase is mediated by compensatory proliferative responses in a lineage-restricted manner (Weng et al., 2018, May et al., 2018). During the chronic phase of injury, however, the progeny of ductal stem cells (marked by K5 or Axin2) partially contribute to the limited number of atypical acini detected in the degenerative gland, suggesting that under some conditions, ductal stem cells can exhibit plasticity and expand their differentiation capacity to produce acinar cells (Weng et al., 2018).

Contrary to irradiation, ligation of the major excretory duct results in almost a complete loss of acinar cells within a few days (Takahashi et al., 2000, Martinez et al., 1982). Removal of the ligature induces a robust regenerative response which begins with emergence of embryonic-like tubular structures followed by regeneration of acini from these structures (Takahashi et al., 2004b, Cotroneo et al., 2008). However, genetic lineage tracing of ductal stem cells using K5-or Axin2-Cre drivers have shown that regeneration of acini in this injury model is independent of ductal stem cells (Weng et al., 2018). The lack of plasticity of ductal stem cells was attributed to the survival of a substantial number of acinar cells and their compensatory proliferation after the obstruction is removed (Aure et al., 2015, Weng et al., 2018). In this model of injury, the severity and extent of acinar cell damage appears to be dependent on the degree by which the duct– associated blood and nerve supplies are obstructed by the ligature (Denny et al., 1997). However, despite the severity of injury, acinar regeneration proceed rapidly, suggesting contribution from non-acinar cells (Walker and Gobe, 1987).

Here to study the cellular mechanisms contributing to acini regeneration after a severe glandular injury, we performed inducible genetic lineage tracing of diverse cell populations within the secretory complex during the injury and recovery of acinar cells using the unilateral duct ligation model. We included the periductal tissues within the ligature to induce extensive acinar cell death, rather than atrophy (Walker and Gobe, 1987). Our analysis revealed a remarkable cellular plasticity of not only K14^+^ ductal stem cells but also more committed epithelial cell types including myoepithelial and intercalated duct cells.

## RESULTS

### Regeneration of acini after severe injury is initiated from K14-expressing progenitors

To assess the extent of injury and dynamics of recovery of acini in a model of severe and reversible injury, the adult FVB mice (8 weeks old, females) were subjected to unilateral ligation of Wharton’s duct and periductal tissues for 3 days; the ligature was removed and both SMGs were collected either immediately (time 0) or at 5, 10 and 20 days after de-ligation for tissue processing and analysis (Fig. 1B). SMGs collected from non-injured mice were used as controls. Histological analysis of SMGs at the time of de-ligation demonstrated an acute loss of acini and GDs with only a few acini remaining at the periphery of lobules, indicating a severe damage to the secretory complex within 3 days of ligation (Fig. 1B and Fig. S1A). The removal of ligature induced a progressive regenerative response, highlighted by rapid formation of tubular structures in the lobules which proceeded by slow reappearance of acini-like structures and subsequent transition into a nearly normal histology by 20 days post-deligation (Fig. 1B and Fig. S1A). However, the gland weight remained at about 76% + 13% (n=3) of normal glands which is consistent with a previous report indicating a smaller size of the injured glands even after 8 weeks of recovery (Cotroneo et al., 2010). Immunofluorescent staining of the control and injured SMGs for aquaporin 5 (Aqp5), a water channel specific to acinar cells (Raina et al., 1995), verified a nearly complete loss of Aqp5 at the time of de-ligation (Fig. S1B), followed by a gradual reappearance of small Aqp5^+^ cell clusters that were often associated with the tubular structures (Fig. 1C, arrow and Fig. S1B). The number of Aqp5-expressing cells returned close to the pre-injury level by day 20 post-deligation (Fig. 1C and Fig. S1B). Evaluation of Ki67 proliferation marker showed a significant increase in the number of proliferating cells at 5 and 10 days after deligation (Fig. 1D), consistent with the rapid expansion of tubular and acini-like structures. Overall, these data validated a model of severe but reversible glandular injury in the adult SMG. It is worth noting that a similar regenerative response was observed when ligature was remove after 7 days (Fig. S1C), however, the ligation period was limited to 3 days throughout the remaining studies for consistency.

The robust proliferative response in tubular structures suggested progenitor cell activity. As K14 marks the only known stem/progenitor cell population in the adult SMGs (Kwak et al., 2016), we assessed the expression of K14 at day 5 and 10 post-ligation when tubular structures dominate the tissue. Analysis of SMG sections stained for K14 revealed a 4- to 5-fold increase in the number of K14-expressing cells (Fig. 1E, Graph). Moreover, these cells displayed a cuboidal morphology and localized mostly to the basal layer of tubular structures (Fig. 1E). To determine if K14^+^ cells directly contribute to acinar regeneration, we performed a short-term pulse-chase experiment in *K14^TRE^:tetOH2BGFP* mouse model which we have previously used to study proliferation dynamics of epithelial cells in the SMG (Kwak and Ghazizadeh, 2015). In this model, administration of doxycycline (Doxey et al.) induces temporary expression of Histone2B-GFP (H2BGFP) fusion protein in K14-expressing cells. Incorporation of H2BGFP in the nucleosome of K14^+^ cells allows monitoring of cell division as nuclear GFP is passed on to their progenies, albeit at reduced fluorescent intensities (Tumbar et al., 2004). Adult bi-transgenic mice (8-10 weeks old females) were subjected to ligature-induced injury and Dox was administered from 3-10 days post-deligation in order to induce label when K14 is broadly expressed in tubular structures (Fig. 1F). The non-injured contralateral gland was used as control. Fluorescent microscopy analysis of SMGs collected at 10 days post-deligation showed robust expression of nuclear H2BGFP in tubular structures (Fig. 1G and Fig. S1D). Immunostaining for K14 protein showed colocalization of nuclear GFP and K14 in control glands, confirming selective expression of H2BGFP by K14-expressing cells (Fig. S1E). In the regenerated gland, however, nuclear GFP was detected not only in the K14^+^ cells but also in other cells within the tubular structures devoid of K14 protein (Fig. 1G, arrowheads), which indicated active cycling of K14-expressing cells and the passage of H2BGFP to their K14^neg^ progeny. Interestingly, immunostaining for Aqp5^+^ showed low-intensity nuclear GFP in a number of newly formed clusters of Aqp5^+^ cells, suggesting that they originated from the H2BGFP-expressing K14^+^ cells (Fig. 1H, arrows). Overall, these data demonstrated that K14-expressing cells in the transitory tubular structures function as progenitor cells for a portion of the newly regenerated acini in this injury model.

### K14^+^ cells in the adult mouse SMG survive and contribute to acinar regeneration after severe injury

The broad expression of K14 in regenerative glands could be either due to expansion of the surviving K14^+^ cells or upregulation of K14 expression in other cell types. To distinguish between these two possibilities, an inducible genetic lineage tracing approach in *K14Cre^TRE^:R26RYFP* mouse model (hereafter referred to as K14-YFP) was used to permanently and selectively mark K14^+^ cells before the injury and map their fate after recovery. We have previously shown that Dox administration for 4 consecutive days in these mice results in yellow fluorescent protein (YFP) labeling of K14-expressing cells, including ductal stem cells and MECs, with an efficiency of about 50% (Kwak et al., 2016). A similar strategy was used to label cells in the adult K14-YFP mice (8-10 weeks old) and 4 days later, mice were subjected to unilateral ligature-induced injury and allowed to recover for 4 weeks before both SMGs were collected for lineage tracing analysis (Fig. 2A). For every mouse, the contralateral gland served as a control to assess the labeling efficiency and YFP expression in the absence of severe injury. In control SMG sections co-stained for K14 and YFP, 49%+3.4% (mean+SD, n=3 glands, >150 cells/gland) of K14-expressing cells were labeled with YFP. Furthermore, after a 5-wk chase period, the progeny of YFP-labeled K14^+^ cells formed small cell clusters in the GD (Fig. 2B), consistent with our previous lineage tracing studies in unperturbed glands (Kwak et al., 2016). Remarkably, analysis of the regenerated SMG revealed vast expansion of YFP-labeled cells into large cell clusters often encompassing the entire secretory complex (Fig. 2C-D). Fate mapping analysis using phenotypic markers demonstrated that in addition to the K14^+^ stem cells and their expected progeny of K19^+^ cells in the GD (Fig. S2A-B), numerous acini (Aqp5^+^) and their associated IDs (cKit^+^) were also expressing YFP (Fig. 2D-F), suggesting multilineage contribution of the initially labeled K14-YFP cells to glandular regeneration after injury. Quantitative analysis showed that all lineage-labeled acini were uniformly labeled with YFP and the majority (92+5%, n=181 acini) were contiguous with YFP-labeled cKit^+^ ID cells (Fig. 2D-F and Fig. S2C). The latter cells, however, did not always expand into the ID/GD junction where ductal stem cells reside (Fig.2F and Fig. S2C). This suggests that some of the lineage-labeled acinar and ID cells are not the progeny of ductal stem cells, instead they might have been derived from MECs, which were also labeled with YFP prior to the injury. However, it was difficult to establish a lineal relationship between MECs and acini in this mouse model as solitary YFP^neg^ MECs were often found surrounding YFP-labeled acini and YFP^+^MECs were surrounding the YFP^neg^ acini (Fig. 2G and S2C, arrows), which is likely due to migratory nature of MECs and sporadic distribution of the initially labeled MECs.

**Figure 2.**
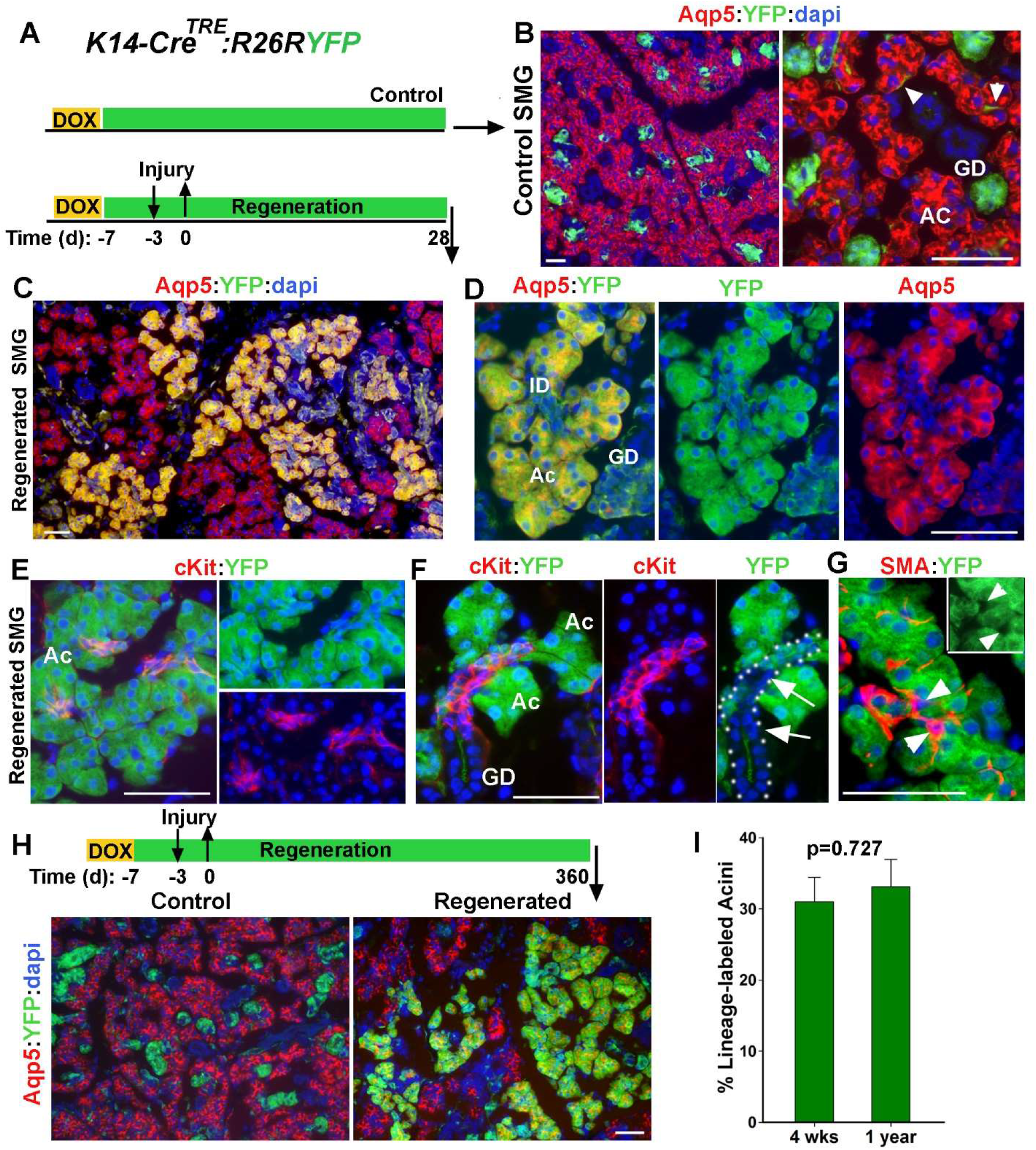
Lineage tracing of K14^+^ cells during injury and recovery of SMG. (A)Time-line of lineage tracing of K14-expressing cells during injury in transgenic *K14Cre^TRE^;R26RYFP* female mice. (B) YFP expression in control SMG stained for Aqp5. Arrowheads point to the YFP-labeled MECs. (C-F) Distribution of YFP-labeled cells in regenerated SMGs co-stained for YFP and Aqp5 or cKit, Dotted line in F outlines ID and GD with arrows pointing to YFP^neg^ ID cells; (G) Distribution of YFP^-^ SMA^+^ cells in labeled cell clusters. Arrowheads point to YFP^neg^SMA^+^. (H) Images of lineage traced cells in control and regenerated gland at 1 year after injury stained for Aqp5. Ac is acini, ID, intercalated duct, GD is granular duct; Scale bars=50 μm. (I) Percentage of YFP-labeled Aqp5^+^ in regenerated SMG collected at 4 weeks or one year after injury. Data are mean+s.e.m (n=3 glands), two-tailed t-test.

Quantification of Aqp5^+^YFP^+^/total Aqp^+^ cells indicated that at 4 weeks post-injury, the proportion of YFP-labeled acini in the regenerated SMGs was about 32.5+3% (mean+SD, n=3 glands). Given the initial labeling efficiency of about 50% for K14^+^ cells, this implies that nearly 65% of the regenerated acini were derived from the surviving K14-expressing cells. Previous studies in skin epithelia have suggested that lineage reversion by committed progenitors is not sustained long-term (Mascre et al., 2012). To determine if lineage-labeled acini derived from K14^+^ cells persist over a long period of time, the chase period was extended to one-year post-injury (Fig. 2H). Remarkably, there was no significant change in the pattern of YFP-labeled cell clusters or the frequency of YFP-labeled Aqp5^+^ cells from 4-wks to 1-year post-injury (Fig. 2H-I), demonstrating persistence of multi-lineage YFP-labeled cell clusters in the regenerated gland. Taken together, these data indicate that K14-expressing cells in the adult SMG survive the severe injury, expand their differentiation potential and contribute to the repair and regeneration of multiple cell lineages in the secretory complex.

### Myoepithelial cells are the primary K14^+^ cell population contributing to acinar regeneration after injury

In the K14-YFP mouse model, both ductal stem cells and MECs were labeled prior to the injury and could have potentially contributed to *de novo* formation of acini after injury. To determine whether MECs contribute to acinar regeneration after injury, we used *αSMACre^ERT2^;R26R-tdTomato* (hereafter referred to as SMA-TdT) in which tamoxifen (TAM)-inducible Cre expression is driven by *Acta2* promoter, to induce specific and permanent expression of the red fluorescence protein tdTomato (TdT) in MECs (Grcevic et al., 2012). Since MECs do not contribute to any other cell lineages during SMG development and throughout adulthood (May et al., 2018), TAM was administered at least 4 weeks before injury to prevent any residual drug from acting in response to injury. Adult transgenic mice (8-10 wks old) were subjected to unilateral duct ligation/deligation and both SMGs were collected at 4 weeks post injury for lineage tracing analysis (Fig. 3A). Immunostaining of sections from control (contralateral) SMGs for cell-specific biomarkers including SMA, K14, cKit and Aqp5 verified exclusive expression of TdT by SMA^+^ cells with an efficiency of 50.5%+4.8% (mean+SD, n=3 glands)(Fig. 3B and Fig. S3A). On the other hand, TdT-labeled cells in the regenerated SMG had expanded into large cell clusters composed of Aqp5^+^ and cKit^+^ cells (Fig. 3B-C and Fig. S3B-D), indicating contribution of the initially labeled MECs to regeneration of both acinar and ID cell lineages likely through the process of transdifferentiation. Interestingly, TdT-labeled acini and ducts were often surrounded by MECs that did not express TdT (Fig. 3D, arrowheads), suggesting a different mechanism for regeneration of MECs than that used for acinar and ductal cells. More importantly, TdT expression was limited to acini and their adjacent portion of the ID but did not advance further toward K14^+^ ductal stem cells (Fig. 3E, arrows and Fig. S3E). Quantification of TdT^+^Aqp5^+^ cells indicated that 24.4+1.1% (mean+SD, n=3 glands) of all acini in the regenerated glands derived from the initially-labeled MECs. The proportion of lineage-labeled acini in SMA-TdT mice was comparable to that in K14-YFP mice (Fig. 3F), identifying MECs are the primary source of regenerated acini in this model of injury.

**Figure 3.**
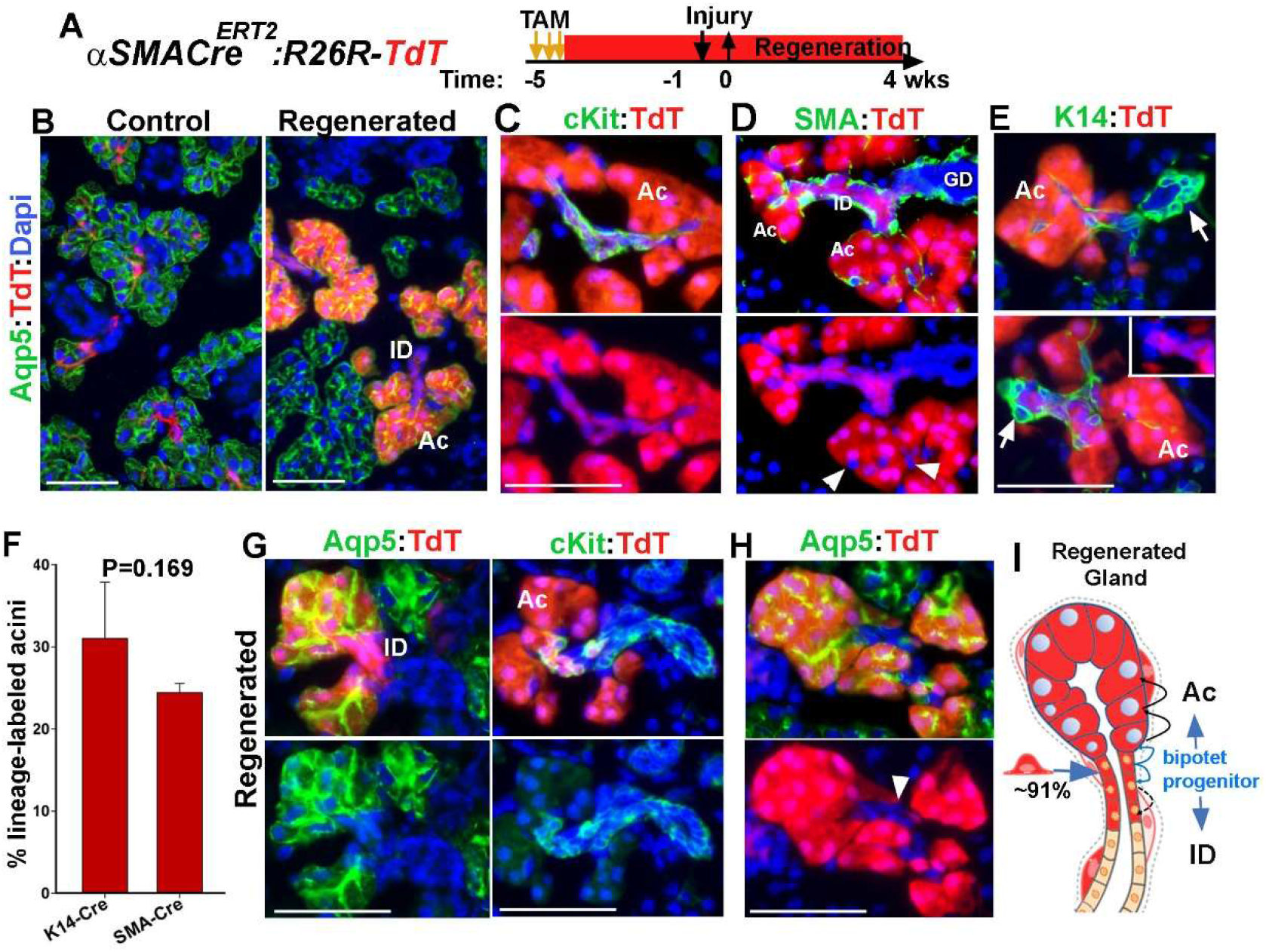
Contribution of Myoepithelial cells to glandular regeneration after injury. (A) Time-line of lineage tracing of MECs with Tamoxifen (TAM) and injury in *αSMACre^ERT2^;R26RTdT* mice. (B) Expression of TdT in control and regenerated SMGs stained for Aqp5. (C-D) TdT distribution in regenerated SMGs stained for cKit, SMA. Green channel is not shown in lower panels to clearly show TdT^neg^ myoepithelial cells (arrowheads) surrounding the lineage-labeled acini. (E) Images of regenerated gland stained for K14. Arrows note the location of ductal stem cells. (F) Quantification of TdT-labeled Aqp5^+^ in the regenerated glands of K14-YFP and SMA-TdT mice at 4-wks post injury, mean+s.d. (n=3 females), two-tailed t-test. (G-H) TdT expression in acini and their associated ID. Sections stained for Aqp5 or cKit and arrowhead points to TdT^+^ acini connected to unlabeled IDs. ID, intercalated ducts; Ac, acini; GD, granular ducts. Blue nuclear staining is dapi. Scale bars = 50 μm. (I) Schematics showing the pattern of lineage labeled cells at 91% of AC/ID junctions and suggesting mechanism. Blue arrows indicate dedifferentiation of MECs to bipotent ductal progenitor and redifferentiation to proacinar and cKit^+^ ID cells. Black arrows indicate self-duplication.

Transdifferentiation is referred to a change in cellular identity from one differentiated cell type to an alternative differentiated state. This may result either from dedifferentiation to a progenitor state followed by redifferentiation to a distinct cell type, or by direct conversion from one cell fate to another (Merrell and Stanger, 2016). To obtain insights into the mechanism of plasticity of MECs, the pattern of TdT expression in the acinar and ID cells within the regenerated SMGs of SMA-TdT mice were analyzed. We found that all acinar cells within the lineage-labeled acini (n=478 from 3 glands) were uniformly expressing TdT suggesting their clonal origin (Fig. 3 and Fig. S3). Analysis of acini/ID junctions showed that although a small number of TdT-labeled acini were connected to unlabeled IDs (Fig. 3H and Fig. S3E), the vast majority (91.1%+5.6%; mean+SD; n=3 gland, a total of 158 acini/ID junctions) were contiguous with at least a few TdT^+^cKit^+^ cell in the ID (Fig. 3C and G, Fig. S3F). Examination of SMG sections where the entire longitudinal dimension of ID’s was visible, showed that in 61.1%+5% of the ID’s (mean+SD, n=3 glands, a total of 97 ID’s), the lineage-labeled cKit^+^ cells did not span the entire length of the duct (Fig. 3G and S3F). This was comparable to the distribution pattern of some of the lineage-labeled acini and ID’s in the regenerated gland of K14-YFP mice in which YFP-labeled cells did not expand toward the position of ductal stem cells (Fig. 2F, arrows and Fig. S2C). The contiguity of lineage-labeled acini and their associated IDs suggests that injury induces dedifferentiation of the initially labeled MEC into a bi-potent progenitor cell which re-differentiate to gives rise to both cKit^+^ ductal cells and proacinar/acinar cells; the latter subsequently self-replicate to repopulate an acinus (Fig. 3I).

### SMG myoepithelial cells have a limited proliferative capacity in culture

The robust response of MECs to injury could be attributed to activation of slowly-dividing stem cells rather than trans- or de-differentiation. In the mammary gland, MECs are maintained as self-sustained cell population but display functional properties of stem/progenitor cells when placed at clonal density in culture (Prater et al., 2014). However, whether MECs display similar characteristics have not been determined. To address this, we first sought to compare the colony forming ability of MECs and ductal stem cells purified from the same gland in adherent cultures. Therefore, we genetically labeled K14-expressing cells in *K14^TRE^:H2BGFP* mice (6 wks old) and then differentially isolated MECs and ductal stem cells based on the expression levels of integrin α6 by FACS (Fig. 4A)(Ogawa, 2003, Lu et al., 2012). Cytospin and immunofluorescent staining of the sorted cell populations for K14 and SMA verified enrichment of SMA^+^ cells with a star-shaped morphology in the α6^high^GFP^+^ cell fraction and those of SMA^neg^ cuboidal-shaped cells in the α6^med^GFP^+^ cell fraction (Fig. 4B). When sorted cells were cultured at clonal density and grown for 2 weeks, ductal stem cells formed large colonies (>0.5 mm in diameter) with a colony forming efficiency of 4.5%+0.36% (Fig. 4C) consistent with their function as tissue stem cells. On the contrary, the colony forming efficiency of MECs was at 0.15%+0.1% as the majority of colonies formed by MECs were abortive (Fig. 4C), indicative of their low proliferative capacity. In addition to clonogenicity, stem cells are assessed by their ability to form organoids in culture (Kopp et al., 2016). When salivary gland cell suspensions are placed in three-dimensional cultures, a small subset of cells form organoids (Lombaert et al., 2008). More recent studies have shown that organoids originate from K14-expressing cells, although no distinction was made between ductal stem cells and MECs (Kwak et al., 2018). To assess the capacity of MECs to form organoids, SMG cells were isolated from TAM-induced SMA:TdT mice (6-8 wks old) and the fate of TdT-labeled MECs were traced in organoid cultures (Fig. 4D). SMG cells from Dox-induced *K14Cre^TRE^:R26RTdT* mice (7-8 wks old; K14-TdT) were used as control. Interestingly, despite a comparable TdT-labeling efficiency in these two transgenic lines (49.5%+4.7% for SMA-TdT and 47%+1.9% for K14-TdT, n=4 mice), traced MECs formed very few TdT^+^ organoids (3.2%+1.1%, n=4 glands, >1100 spheres/gland) when compared to traced K14-expressing cells (47.6% +7.6%, mean+SD, n=4 glands, >850 spheres/gland) (Fig. 4E-F). The direct correlation between the TdT-labeling efficiency of K14-expressing cells *in vivo* and that of organoids in culture indicates that almost all organoids originate from K14^+^ ductal stem cells. Overall, functional analysis of MECs in two culture systems indicate that salivary gland MECs have low proliferative capacity in culture and may not function as stem cells. Therefore, the robust regenerative response of MECs to injury is likely due to transdifferentiation of MECs rather than stem cell activation.

**Figure 4.**
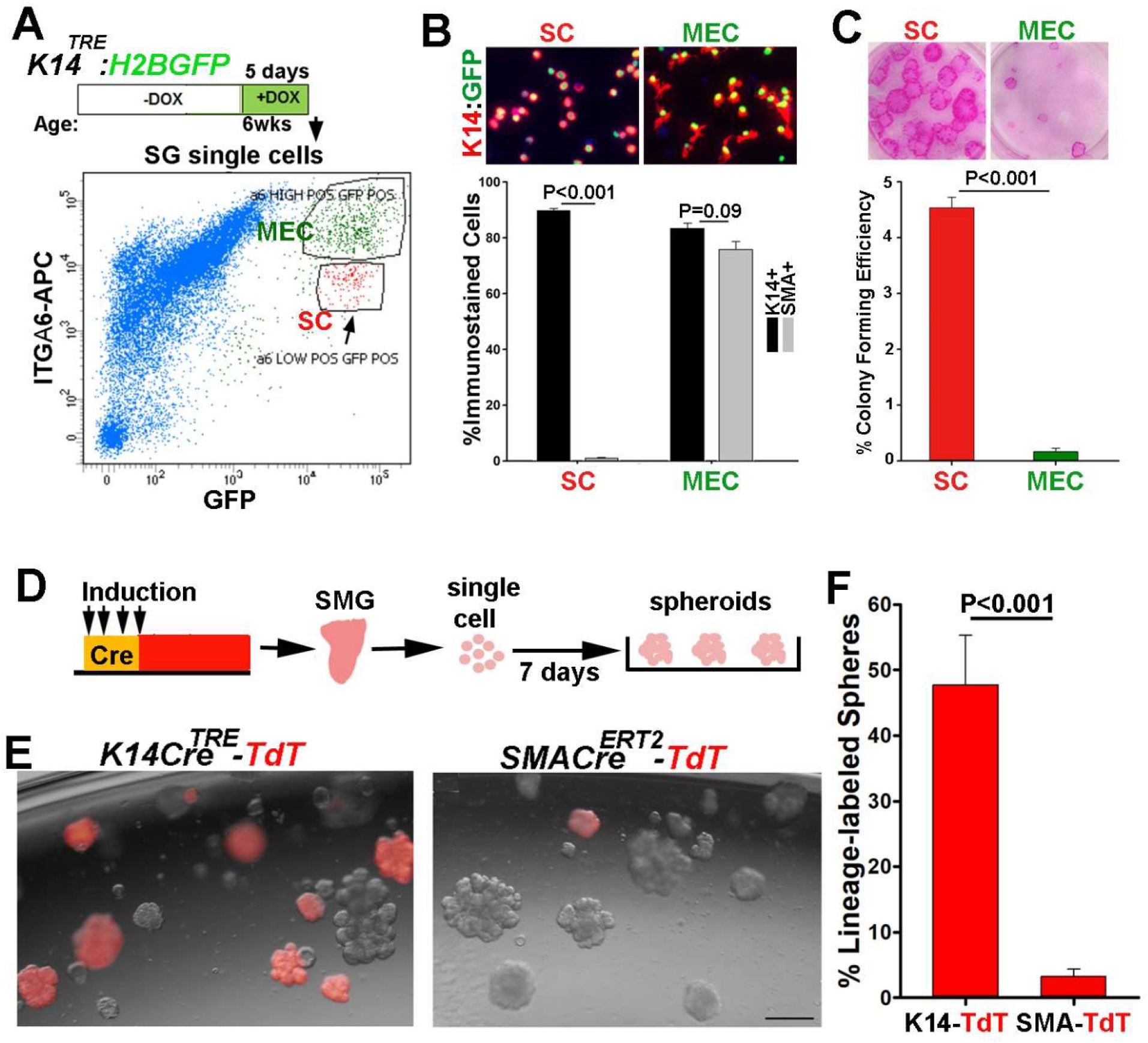
MECs do not contain highly proliferative progenitors. (A) FACS profile of Dox-induced K14-H2BGFP labeled cells stained with APC-anti-integrin α6 antibody. Arrow points to the ductal stem cell population expressing lower levels of integrin-α6. (B) Cytospin analysis of sorted GFP^+^α6^hi^ (MEC) and GFP^+^α6^med^ (SC) cells. Graph shows percentages of K14^+^ (black bars) and SMA^+^ (gray bars) cells in each population (n=200 cells/population). (C) Colony formed from sorted cells seeded at 500 cell/well, grown for 2 weeks and stained with Rhodamine. Graph is colony forming efficiency (diameter > 0.5 mm). (D) Strategy used to induce label in either SMA^+^ or K14^+^ cells *in vivo* and analyze their fate in organoid cultures. (E) Images of organoids established from SMG cells of K14-TdT and SMA-TdT mice after 7 days in culture. Scale bars=200 μm.(F) Percentage of TdT-labeled organoids in culture (>1100 organoids counted/gland). Values are mean+s.e.m, from at least two independent experiments. two-tailed t-test, p is indicated in the figure.

### Ductal stem cells undergo a unipotency to multipotency switch in response to injury

To determine if K14^+^ ductal stem cells make any contribution to acinar regeneration in this injury model, we traced the lineage of ductal stem cells in *Axin2Cre^ERT2^;R26RTdT* (Axin2-TdT) mice (Fig. 5A). Axin2Cre driver has been shown to target a subset of K5^+^/K14^+^ ductal cells but is not active in MECs (Weng et al., 2018). In these experiments, K14-TdT mice were treated in parallel and used for comparative analysis. In the control SMG of Axin2-TdT mice, TdT-labeled cells were localized to small cell clusters near the ID/GD junctions which consisted of K14^+^ ductal stem cells and their progeny (Fig. 5B, arrows), but excluded cells expressing cKit, Aqp5 or SMA (Fig. 5B). Moreover, the size and distribution frequency of TdT-labeled cell clusters in Axin2-TdT mice were comparable to those in K14-TdT mice, with the exception that TdT-labeled MECs appeared only in the latter (Fig. 5C and S4A-B). Interestingly, analysis of TdT expression in the regenerated SMGs showed a few but relatively large TdT-labeled cell clusters composed of ductal and acinar cells (Fig.5D-F and Fig. S4B). In these cell clusters, TdT could be traced from K14^+^ ductal cells (Fig. 5D, arrow) along the length of ID (cKit^+^) into acini (Fig. 5D-E), indicating multi-lineage contribution from ductal stem cells. However, SMA^+^ cells (or K14^+^ basal cells) juxtaposed to the TdT-labeled cell clusters were completely clear of TdT expression (Fig. 5D, arrowheads and Fig. S4D, arrows), supporting the independence of MEC lineage from Axin2^+^K14^+^ ductal stem cells. Quantification of TdT^+^Aqp5^+^ cells showed a significantly lower frequency of lineage-labeled acini in the regenerated SMGs of Axin2-TdT mice (4.1%+0.5%) when compared to those in K14-TdT mice (28%+5%) (Fig. 5G and Fig.S4B). This is consistent with the dominant contribution of MECs to acinar regeneration in this model of injury. Taken together, these data indicate that injury does induce multilineage differentiation capacity of ductal stem cells (Fig. 5H), however, their contribution to acinar regeneration is not significant as MECs, possibly due to their scarcity and physical distance from acini.

**Figure 5.**
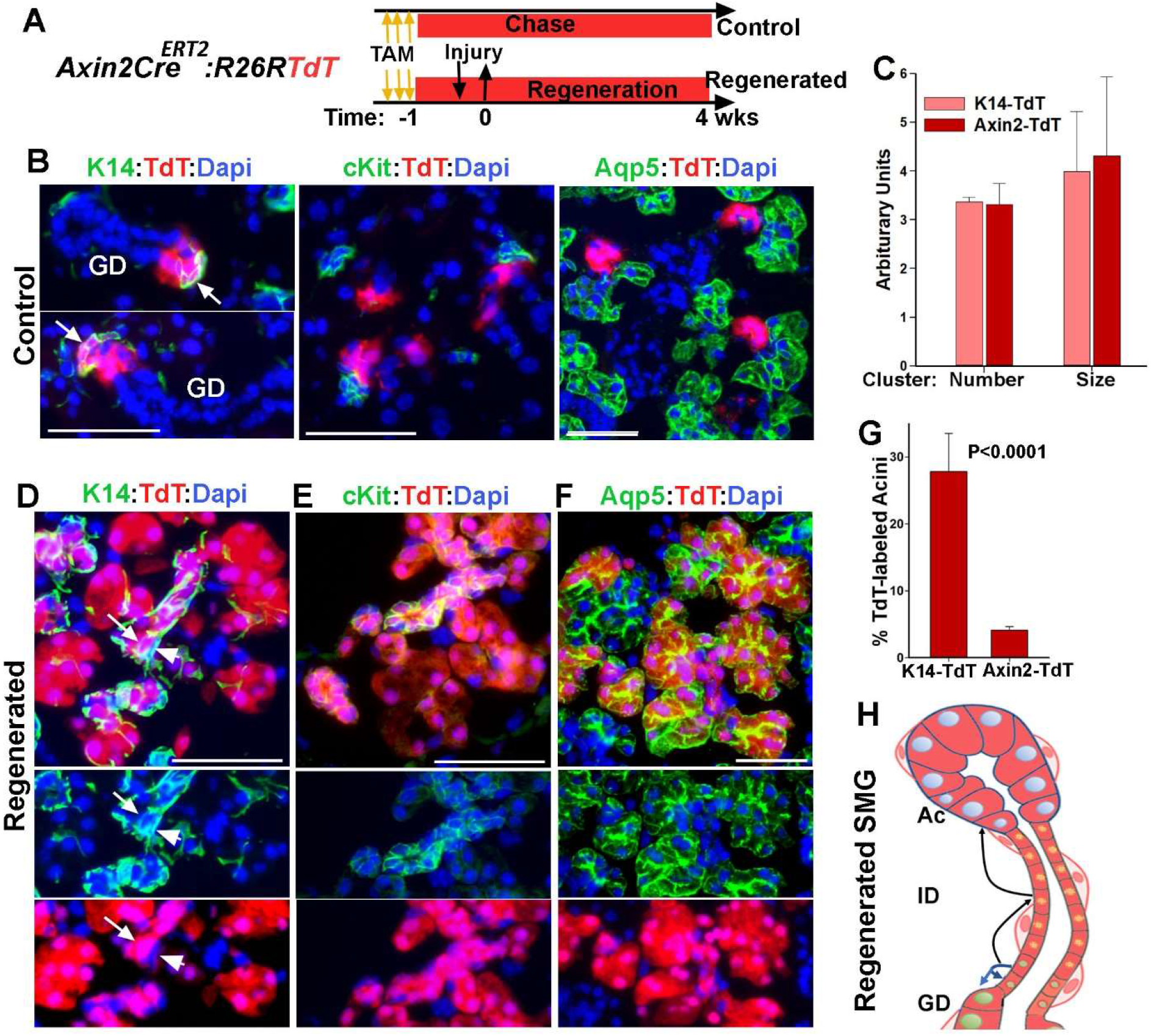
Severe injury induces a switch to multipotency in ductal stem cells. (A) Time-line of lineage tracing of ductal stem cells in *Axin2-Cre^ERT2^;R26RTdT* in control and injured glands. (B) TdT expression in control SMG stained for K14, cKit and Aqp5. Arrows notes K14^+^ ductal stem cells. (C) Comparative analysis of TdT-labeled clusters in control SMGs of Axin2-TdT and K14-TdT mice after a 5-wk chase period. (D-F) TdT expression in the regenerated SMG stained for K14, cKit and Aqp5. Arrow and arrowheads in D notes ductal stem cells and MECs, respectively. Nuclear blue is dapi. Scale Bars=50 μm. (G) The percentage of TdT-labeling in acinar cells in the regenerated glands K14-TdT and Axin2-TdT mice. Data are presented as mean+s.d. (n=3 females), two-tailed t-test. (H) Summary of results of lineage tracing of ductal stem cells. Blue arrows show normal fate of stem cells and black arrows show expansion of their differentiation potential.

### cKit^+^ ductal cells function as reserved progenitors for acinar cells

Our results, so far, indicated that more than half of newly formed acini originate from MECs and ductal stem cells, however, the origin of the remaining acini is unclear. The injury-induced plasticity of MECs toward acinar cell differentiation appeared to transition through cKit^+^ bipotent progenitors as an intermediate cell population. In the adult SMG, cKit marks the ID cells as well as tuft cells that are scattered in the larger ducts, however neither cells give rise to any other cell types within the gland (Kwak et al., 2018). More importantly, when placed in culture, cKit^+^ cells do not form organoids indicating that they do function as progenitor cells in the adult gland (Kwak et al., 2018). However, during SMG development, cKit^+^ marks progenitors that differentiate into ID and acinar cells (May et al., 2018). Therefore, we hypothesized that severe injury provokes dedifferentiation of cKit^+^ ID cells to an earlier progenitor-like state that will generate acinar cells. To test this, we traced the lineage of cKit^+^ cells during ligature-induced injury and recovery using *cKitCre^ERT2/+^*; *R26RTdT* (cKit-TdT) transgenic line (Fig. 6A). The specificity of TdT expression in cKit^+^ cells were verified in control SMG (Fig. 6B-C and Fig. S5A-B) and the TdT-labeling efficiency of cKit^+^ ID cells was estimated to be at 71%+7.4% (Fig. S5B; mean+SD, n=3 glands) which was consistent with our previous studies (Kwak et al., 2018). Fate mapping of cKit-lineage cells in the regenerated SMGs revealed expansion of the TdT-labeled cells from ID’s into acini (Fig. 6B and D and Fig. S5C). This expansion was directed only toward acini as we did not detect any TdT-expressing K14^+^ ductal cells or MECs in the regenerated gland (Fig. 6E, arrowheads and Fig. S5D-E). These data suggested that in response to injury, c-Kit^+^ ID cells function as reserve cells contributing to recovery of secretory units (Fig. 6G). Quantitative analysis of Aqp5^+^ cells indicated that 26%+8.9% (mean+SD, n=3glands) of acinar cells were expressing TdT. Given the initial labeling efficiency of about 70% for cKit^+^ ID cells, these results indicated that approximately 37% of the total regenerated acini derived from cKit^+^ ID cells.

**Figure 6.**
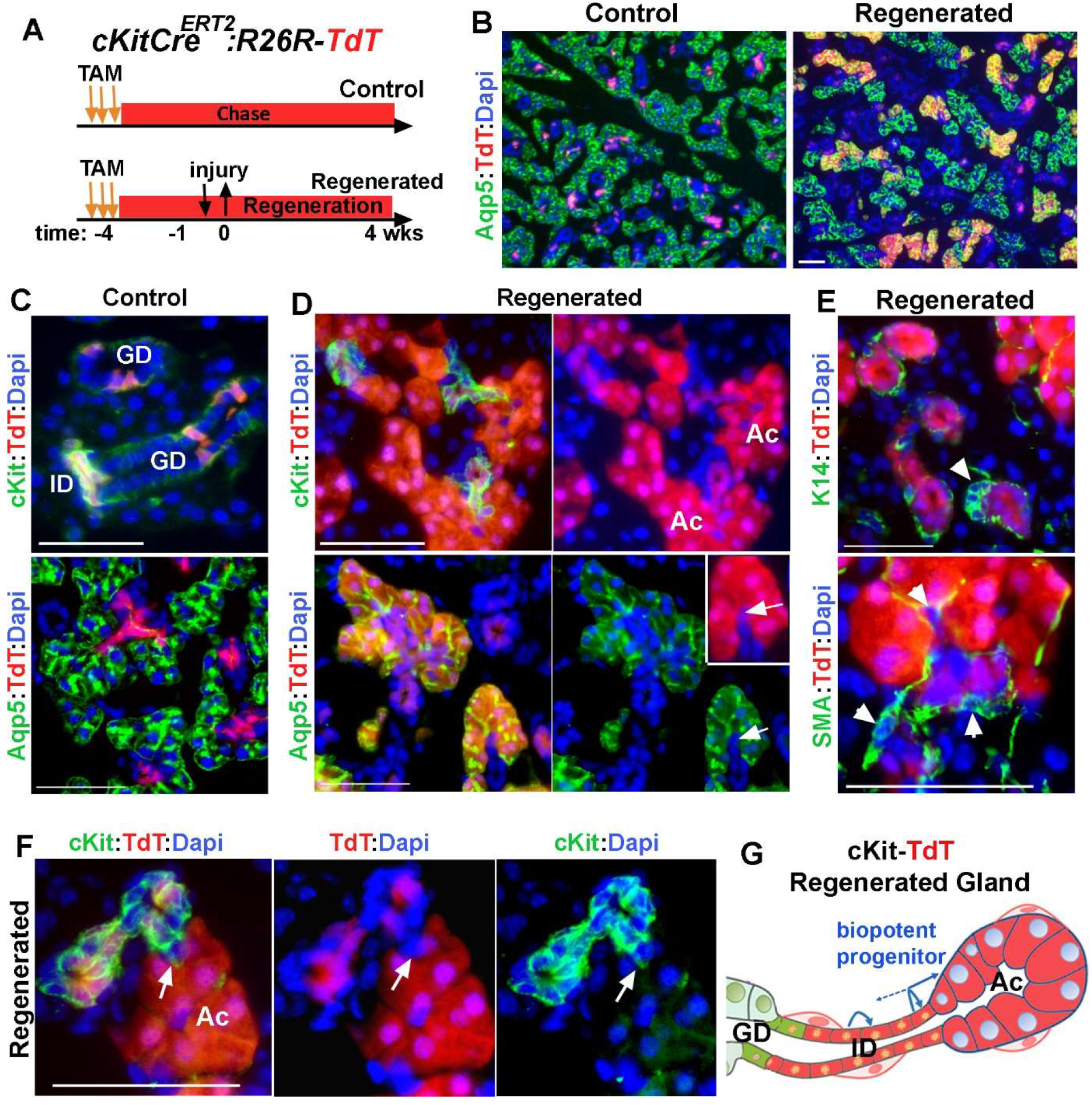
Contribution of cKit^+^ ductal cells to regeneration of acini. (A)Time line of label induction and lineage tracing of c-Kit^+^ cells during injury and regeneration. (B-D) TdT-lineage traced cells in control and regenerated glands stained for Aqp5 or cKit. Arrows in D points to a single TdT^+^ ID cell contiguous with TdT^+^ acini. (E) Images of TdT-labeled cell clusters stained for K14 and SMA. Arrowhead points to ductal stem cells. (F) A small number of TdT-labeled acini are not contiguous with TdT-labeled cKit^+^ cells (arrow). Nuclear blue staining is dapi. ID, intercalated ducts; AC, acini; GD, granular duct. Scale bars=50 μm. Images are representative of 3 glands from female mice. (G) Summary of results of lineage tracing of ID cells indicating dedifferentiation of cKit^+^ cells to a bipotent progenitor cell population which re-differentiated to both ID and acinar cells. Distal ID cells may self-replicate.

TdT expression pattern in regenerated glands of cKit-TdT mice was almost identical to that in SMA-TdT mice. While all acinar cells within an acinus appeared to be clonal in origin (Fig. 6D-F and Fig.S5C-G), TdT expression in the ID was more heterogeneous with about 55% of the ID’s (n=142) composed of a mixture of TdT^+^ and TdT^neg^ patches of cKit^+^ cells (Fig. 6F and Fig. S5G). Moreover, analysis of Acini/ID junctions demonstrated that despite the presence of a few TdT^+^ acini connected to non-labeled ID’s (Fig. 6F, arrow and Fig. S5G), in the majority of lineage labeled acini (91.6%+5.7%, mean+SD; n=100 from 3 glands), TdT could traced from acini to the ID (Fig. 6D and Fig. S5F), indicating that they derived from a cKit^+^ ductal progenitor. This pattern is consistent with dedifferentiation of cKit^+^ cell to an earlier state when cKit^+^ cells function as bi-potent progenitor cells (May et al., 2018). The similarity in organization of lineage-labeled cells in the regenerated glands of cKit-TdT and SMA-TdT mice suggests that both myoepithelial and intercalated ductal cells dedifferentiate into a common bipotent progenitor-state before redifferentiation to proacinar cells (Fig. 7).

**Figure 7.**
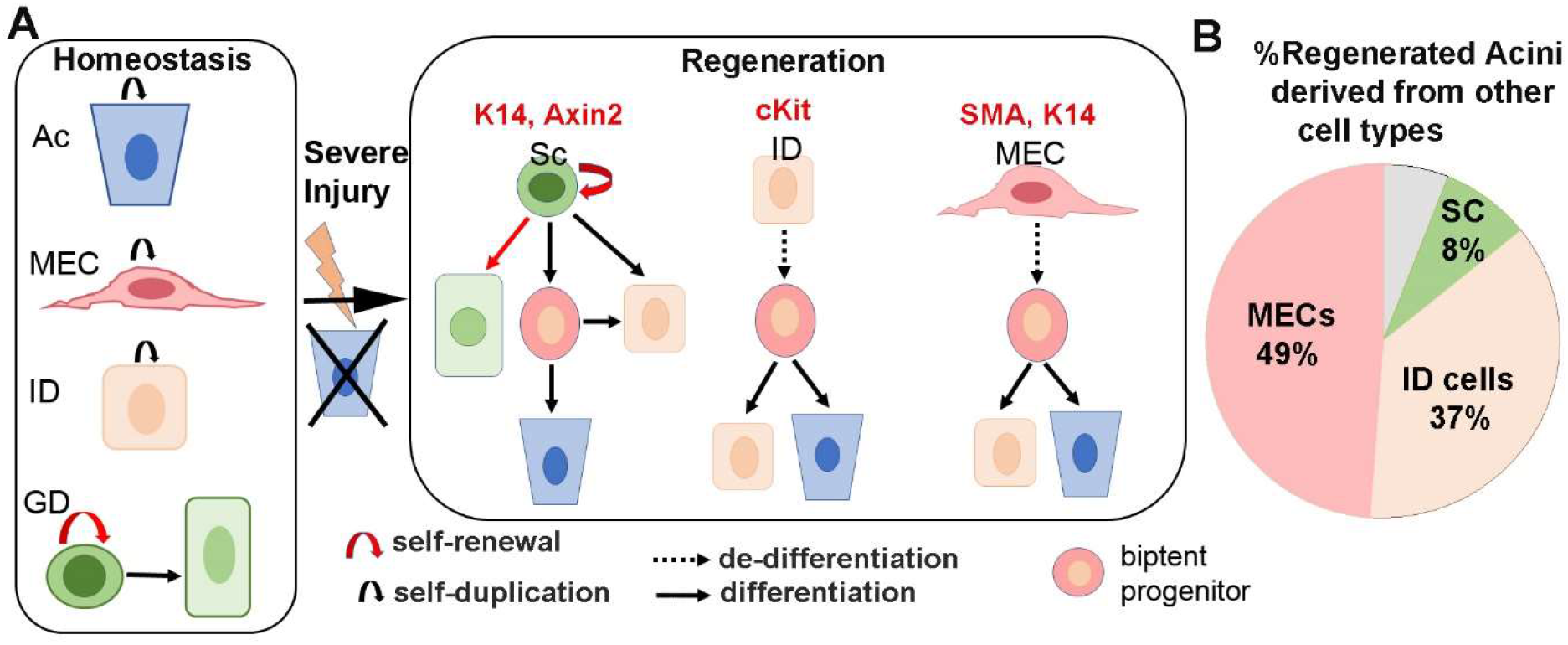
Severe injury provokes lineage plasticity of ductal and myoepithelial cells toward acinar cell differentiation. **(A)** The cell lineages in the adult SMG are maintained in lineage-restricted manner under homeostasis. Ac is acini, MEC is myoepithelial cells, ID is intercalated duct, SC is ductal stem cell and GD is granular duct cells. Severe injury induces multipotency in ductal SC and promote dedifferentiation of cKit and MECs into a bi-potent progenitor-like state which re-differentiates to ID and Ac. The gene derivers used for lineage tracing of each cell lineage is indicated on the top. (B) The contribution of each cell lineage to acinar regeneration after correcting for the initial labeling efficiency.

## DISCUSSION

The robust regenerative responses of salivary glands to injury suggest that there are efficient and effective mechanisms in place to maintain the secretory component and gland function. Here, we used a model of reversible glandular injury in which acinar cells are severely depleted and therefore cannot undergo compensatory self-replication to reveal the cellular mechanisms underlying *de novo* formation of acini in the adult gland. By tracing the lineage of three distinct non-acinar cell populations that reside in the secretory complex we uncovered a remarkable cell plasticity and reversibility in epithelial cells within this compartment. Our findings reveal that both stem cells and more committed cells contribute to regeneration of secretory units following injury to allow rapid recovery of glandular tissue. We show that upon injury, K14^+^/Axin2^+^ ductal stem cells expand their differentiation capacity to include cKit^+^ ductal and acinar cell lineages, suggesting the induction of a unipotency to multipotency switch (Fig. 7). However, this is not the major mechanism by which acini are regenerated. Intriguingly, our findings demonstrate that most of the regenerated acini derive from more committed cells including MECs and cKit^+^ ductal cells. Contrary to ductal stem cells, these cells display a very slow turnover rate and normally do not contribute to any other cell lineages in the adult gland (Kwak et al., 2018, Kwak and Ghazizadeh, 2015, May et al., 2018). However, in response to a severe insult, they participate in tissue repair and regeneration by dedifferentiating into a common bipotent progenitor-like state and subsequently re-differentiate to acinar cells (Fig. 7). Therefore, plasticity of more committed parenchymal cells is the predominant mechanism of acini regeneration in response to a severe injury.

We used ligation/deligation of the main salivary duct to study the degree of cellular plasticity in non-acinar cells in the secretory complex of adult SMGs after a severe injury. Although this injury model has been used extensively to study salivary gland repair and regeneration, the source of acinar regeneration that follows deligation has been a subject of much speculation. While some studies found that acinar regeneration occurred mainly by self-duplication of surviving acinar cells (Burgess and Dardick, 1998, Aure et al., 2015), others suggested that ductal cells differentiate to acinar cells (Cotroneo et al., 2010, Takahashi et al., 1998, Walker and Gobe, 1987). The compelling evidence provided by our fate-mapping analyses of two distinct ductal cell populations (cKit^+^ cells and K14^+^/Axin2^+^ cells) are consistent with the latter and underlines the plasticity of ductal cells toward acinar cell differentiation during the repair of secretory complex. Our data, however, contradicts previous lineage tracing studies that did not find any contribution from K5^+^ or Axin2^+^ ductal cells to regeneration of acini following duct ligation and deligation (Weng et al., 2018). This is likely related to the severity of obstruction-induced injury and the extent of acinar cell damage (i.e., atrophy vs. cell death) which appears to be dependent on the degree to which the duct–associated blood and nerve supplies were obstructed by the ligature (Denny et al., 1997). In our study, the inclusion of periductal tissue within the ligature induced a severe damage to secretory cells within 3 days of ligation, while in a study reported by Weng et al. (Weng et al., 2018), a substantial number of acinar cells were present even after two weeks of duct ligation. Indeed, in a parallel comparison between obstructing salivary duct with and without periductal tissue, we confirmed a lack of contribution from non-acinar cells to regenerated acini in the latter injury model (data not shown). Therefore, substantial damage causing obliteration of acini is required to provoke plasticity and reversibility of other cell types toward differentiation to acinar cells. Clearly, the type of injury and the resulting wound environment induced by obstruction of duct and associated vascular and neural tissues is quite different than those induced by the isolated duct obstruction and further investigation will be needed to elucidate these differences.

Our analyses provide the first direct evidence demonstrating the injury-induced plasticity of MECs to other cell types in salivary glands. Although previous studies have described the presence of proliferating MECS covering newly formed acini during regeneration of submandibular gland after duct deligation (Takahashi et al., 2004a), whether or not these cells play a role in supporting acinar cell differentiation was not known. MECs are found on the basement membrane of all exocrine glands and have been known to display functional properties of stem/progenitor cells in some organs such as mammary gland and submucosal glands of the lung (Tata et al., 2018, Lynch et al., 2018, Prater et al., 2014). Although mammary gland MECs are a long-lived lineage-restricted cell population *in vivo*, when placed in adherent cultures more than half of MECs are clonogenic indicating their robust proliferative potential (Prater et al., 2014). Salivary gland MECs are also a self-sustained cell lineage during development and throughout adulthood (Kwak and Ghazizadeh, 2015, May et al., 2018, Song et al., 2018); yet, their robust contribution to ductal and acinar cells in response to injury suggested that salivary gland MECs may also contain stem cells or progenitors. However, using two well-established culture system that have been used to assess stemness in various tissues, we found no evidence for the existence of highly proliferative progenitors in salivary gland MECs. Given that MECs do not appear to function as progenitors, their multilineage contribution to the regeneration of secretory complex can be best explained by reversion of differentiation into a progenitor/stem cell-like state (Mannik et al., 2010, Blanpain and Fuchs, 2014). The distribution pattern of MEC-derived lineage cells in the regenerated secretory complex supported this conclusion. The uniform fluorescence labeling (YFP or TdT) of cells within regenerated acini suggests that each newly formed acinus originated from a single labeled proacinar cell which was a product of MEC dedifferentiation. Given that the efficiency of Cre recombination in the K14-Cre^TRE^ or SMA-Cre^ERT2^ mice was about 50%, contribution from more than one proacinar would have resulted in a mosaic pattern of labeled and unlabeled acinar cells, at least within some acini. Furthermore, more than 90% of the lineage-labeled acini were contiguous with at least a single lineage-labeled cKit^+^ cell in the associated ID. More importantly, a similar pattern was observed in cKit-lineage-labeled cell clusters suggesting that both cKit^+^ ID cells and MECs dedifferentiate to a common bi-potent progenitor-like state to generate a cKit^+^ ductal cell and a proacinar cell which subsequently self-replicates to repopulate/regenerate the acinus. Interestingly, the presence of transitory tubular structures during the regenerative phase (5-10 days after deligation) of recovery in this model of injury recapitulates the perinatal stage of gland development, both morphologically and chronologically (Fig. 1)(Cotroneo et al., 2010). During SMG development, the differentiation of proacinar, intercalated ductal and myoepithelial cells begins at the same time (E15-E16) from a common K14^+^/cKit^+^/Sox10^+^ progenitor and Sox10 has been identified as key transcription factor controlling plasticity and multipotency of this cell population (Lombaert et al., 2013, Athwal et al., 2019). This supports our conclusion that both cKit^+^ cells and MECs in the injured adult gland revert back to a common progenitor-like state. Interestingly, despite the recently recognized role for Sox10 in the regulation of multipotency during SMG development (Athwal et al., 2019), this transcription factor is broadly expressed in acini, ID and MECs in the unperturbed adult mouse SMG (data not shown) (Ohtomo et al., 2013). What are the major constituents of the embryonic-like phenotype/microenvironment promoting the plasticity of ductal and myoepithelial cells and how Sox10 and many other factors including inflammation may function in this model of injury awaits further investigation.

In summary our findings reveal a remarkable plasticity of ductal and myoepithelial cells toward acinar cell differentiation provoked by a severe injury to secretory complex in the adult salivary gland. Salivary glands rely in recruitment of committed and fully differentiated cell populations as well as stem cells to ensure rapid regeneration and recovery of secretory cells. Further studies in this model of injury will provide greater clarity on environmental signals that control fate decisions in the context of severe glandular injury. Such studies are likely to yield important information regarding epithelial tissue plasticity that could be employed to develop new effective therapies aimed at reversing hyposalivation.

## MATERIALS AND METHODS

### Animal Studies

The care and experimental use of animals including the surgical procedures were approved by the Institutional Animal Care and Use Committee (IACUC protocol#287271) of the Stony Brook University, Stony Brook, New York, USA. Animals were maintained in the Stony Brook University-Health Science Center Division of Laboratory Animal Resources, an AAALAC-accredited center, and housed in groups of up to five mice per cage in a pathogen-free facility on a 12 hr light/dark cycle provided with food and water *ad libitum.* All animal experiments were performed in accordance with institutional guidelines set forth by the State University of New York. Adult mice (>8 weeks of age) of the following transgenic strains were used: Transgenic tetO-H2BGFP (stock# 005104), *K14rtTA* (stock#008099), *Axin2-Cre^ERT2+/-^* (stock#018867) and, Cre reporter lines R26R-EYFP (stock# 006148) and R26R-tdTomato (stock# 007914) were purchased from Jackson Laboratory (Bar Harbor, Maine). *K14Cre^TRE^* and *cKit-Cre^ERT2+/-^* transgenic lines have been described previously (Kwak et al., 2016, Klein et al., 2013). *αSMA-Cre^ERT2+/-^* were described previously and generously provided by Dr. Ivo Kalajzic’s lab (Grcevic et al., 2012). *cKit-Cre^ERT2+/-^* and *αSMA-^CreERT2+/-^* were maintained on a C57BL/6 background. Mice were genotyped by polymerase chain reaction using mouse genomic DNA from tail biopsy specimens. For lineage tracing experiments Cre-mediated recombination was induced in mice by replacing regular diet with doxycycline-(1g/Kg pellet) or tamoxifen-containing diet (250 mg/Kg pellet, BioServ, Flemington, NJ) for 4-5 days. Mice were returned to regular diet for 4 days to 4 weeks before subjected to injury. Surgical procedures were performed under anesthesia and using aseptic techniques. A small incision was made on the lateral side of the neck and the main excretory duct on one side of the neck was dissected under a surgical stereoscope. A small plastic microtube (2 mm in length; sterile intramedic polyethelene tubing, Cat #: 427400, Becton Dickinson) was placed along the duct and tied together using 6-0-black-silk suture (Fisher Scientific), and the incision was closed by surgical sutures. At the time of deligation, mice were anesthetized and external sutures were removed to expose the ligated duct, microtube was removed and suture was cut to release the obstruction. The area was rinsed with saline and the incision was closed. Animals were left to recover for the indicated period of time before both ligated and contralateral glands were harvested and processed.

### Tissue processing and Immunofluorescent Staining

SMGs were harvested and the attached fat and connective tissues were dissected. SMGs were either fixed in 10% formalin for 24 hrs for paraffine embedding or in 4% paraformaldehyde at 4 °C for 1 hour followed by 30% sucrose before cryopreservation. Five-micron cryosections were prepared, dried for at least 2 hrs, rehydrated, and immunostained as described previously (Kwak and Ghazizadeh, 2015). The following primary antibodies were used: anti-GFP/YFP (chicken, 1:2000, Cat#ab13970 Abcam) for the R26R-YFP reporter strain, anti-c-Kit (Rat, 1:100, Cat# 14-1171-82, eBioscience), anti-Aqp5 (Rabbit, 1:100, Cat#178615, EMD Millipore, Billerica, MA), anti K19 (Rat, 1:10, TROMA-III, DHSB, University of Iowa), anti-K14 (Rabbit, 1:2000, Cat# PRB-155P, Covance, Berkeley, CA), anti α-SMA (Rabbit, 1:100, Cat#ab5694, Abcam, Cambridge, MA) and anti-integrin α6 (APC-conjugated, Cat#17-0495-82, eBioscience). Alexa-594 or Alexa-488 conjugated secondary antibodies were from Molecular Probes (Invitrogen, Eugene, OR). TdT was detected by direct fluorescence. Sections were mounted in Vectashield^®^ mounting medium containing 4′,6-diamidino-2-phenylindole (DAPI; Vector Laboratories, Burlingame, CA). Fluorescent staining was visualized using a Nikon E800 fluorescent microscope and images were captured using NIS-Elements (Nikon Instrument Inc., Melville, NY). Image J was used for quantification of labeled cells using images from a pool of at least 5 images taken at 100X or 15 images taken 400X from multiple sections stained for each control and regenerated gland.

### SG Cell Preparation, Cell Sorting and Cultures

SMGs were dissected, dissociated by mechanical and enzymatic digestion (0.025% Collagenase, 0.05% hyaluronidase, 1U/ml dispase) to single cell suspensions as described previously (Kwak and Ghazizadeh, 2015). For adherence cultures, single cell suspensions prepared from H2BGFP-labeled SMGs were re-suspended in PBS-1% bovine serum albumin and stained with APC conjugated antibody to integrin α6 (clone GoH3, from eBioscience) for 20 minutes at 4°C in dark. Cells were washed with PBS-1% BSA and re-suspended in PBS-2% fetal bovine serum. Cell sorting was performed on FACSAria-III with FACSDiva software (BD Bioscience). Sorted cells were plated at 50 cells/cm^2^ in the presence of gamma-irradiated 3T3 fibroblasts as described previously (Kwak and Ghazizadeh, 2015). Colonies were visualized and counted after staining with 1% Rhodamine B (Guercio et al.). For organoid cultures, single cell suspensions of SMGs were re-suspended to 1X10^4^ cells in 40 μl media, mixed with 60 μl Matrigel (BD Biosciences, San Jose, CA) on ice, seeded in the periphery of 12-well tissue culture plate and transferred to 37°C incubator for 20 min to solidify. Gels were covered in a modified salisphere growing media containing epidermal growth factor (20 ng/ml), fibroblast growth factor-2 (20 ng/ml), N2 supplement, insulin (10 μg/ml), dexamethasone (1 μM), 5% fetal bovine serum and 10 μM ROCK inhibitor Y-27632 (Guercio et al.), and cultured for 7 days before analyzed by fluorescent phase-contrast microscopy. Images were captured from the entire periphery of the wells and used to quantify the number and percentage of labeled spheres in culture.

### Quantification and Statistical Analyses

Typically, 3-4 sections taken at least 50 μm apart were analyzed for each animal, and the average values for each animal were used to calculate the mean ± SD. For analysis of lineage-labeled acini, 100X images were used to measure the area of the lineage labeled-Aqp5^+^ cells to total Aqp5^+^ areas expressed in pixel density. For quantification of labeling efficiency in the targeted cell population and fate mapping analysis 400X images were used and the number of labeled cells within a specific population or location were quantified. All statistics were performed using SPW12 software (Systat software Inc., San Jose, CA). Differences among means were evaluated by un-paired 2-sided Student’s t test when comparing two groups and one-way analysis of variance and the Tukey’s HSD post hoc comparison when comparing several groups. Significance was set at p<0.05.

## Acknowledgments

This work was supported by the NIH-NIDCR (R21DE022959 to S.G.). We thank Dr. Lucille London for critically reviewing this manuscript, Dr. Ivo Kalajzic for donating the aSMACre^ERT2^ mice and Laurie Levine for technical assistance with our mouse colonies.

